# Important effect of the X chromosome in genomic evaluations of reproductive traits in beef cattle

**DOI:** 10.1101/2020.06.30.178558

**Authors:** Iara Del Pilar Solar Diaz, Gregório Miguel Ferreira de Camargo, Valdecy Aparecida Rocha da Cruz, Isis da Costa Hermisdorff, Caio Victor Damasceno Carvalho, Lucia Galvão de Albuquerque, Raphael Bermal Costa

## Abstract

The aim of this study was to evaluate the efficiency of inclusion and the prediction ability of the X chromosome for reproductive (occurrence of early pregnancy – P16 and age at first calving - AFC) and andrological traits (scrotal circumference -SC) in a herd of Nellore beef cattle herd. 3,263 genotypes of females and males were used. Genomic prediction for SC, AFC and P16 was carried out considering two scenarios: 1) only autosomal markers or 2) autosomal + X chromosome markers. To evaluate the effect of inclusion of the X chromosome on selection, the responses to the selection performed were compared including or not the X chromosome in the evaluation of the traits. Higher heritability estimates were obtained for SC (0.40 and 0.31), AFC (0.11 and 0.09) and P16 (0.43 and 0.38) for the analyses including the X chromosome compared to those without. The percent reduction on mean genomic breeding values when selection was based on the results of analysis that did not include the X chromosome to 1, 5 and 10% of the top males, was for SC slightly more than 7% of the mean genomic breeding value of the selected animals. For P16, the loss can reach more than 4%, while this loss does not seem to be as important for AFC. Average predictive correlation of 0.79, 0.98 and 0.84 for SC, AFC and P16 was obtained, respectively. These estimates demonstrate that inclusion of the X chromosome in the analysis can improve the prediction of genomic breeding values, especially for SC.

## 1. Introduction

Genomic selection is important for the evaluation of reproductive traits. These sets of traits are difficult to assess and to measure directly and generally show low additive genetic variability. This low additive genetic participation in the expression of the phenotype can be explained by the low adaptive value as a result of selection for growth traits (Jensen and Andersson, 2005; Katz, 2008). Part of it is also explained by the management imposed by breeders (such as weight and age limits to enter breeding) and by the lack of measurement of phenotypes in low-performance animals.

Marker effects are usually estimated in evaluations that consider only the effects of autosomes. The sex chromosomes are excluded because of the mechanism of dosage compensation of X chromosome-linked genes in females and because males are hemizygous for the X chromosome (Couldrey et al., 2017). On the other hand, the X chromosome is the second largest chromosome of the genome and therefore harbors many genes that can affect different phenotypes (Berry et al., 2017). When X chromosome markers are disregarded, the direct and epistatic effects of its genes are not measured and the effects of autosomal markers may be overestimated.

According to Ferreira and Franco (2011), considering the effects of sex chromosomes in the field of animal reproduction is essential because a large part of the mechanism involved in normal embryonic development comes from how the X chromosome is established in the population. Furthermore, both X chromosomes are active in the ovaries for gamete production (Arnold et al., 2016), i.e., there is no inactivation of one of the X chromosomes in this organ as occurs in other organs, further emphasizing the participation of this sex chromosome in reproductive events. Complementarily, the autosomes harbor genes with diverse functions and extremely heterogeneous expression patterns, while the sex chromosomes are rich in genes related to development and sexual reproduction (Liu, 2019). Within this context, some studies reported interesting results regarding the roles of the X chromosome in reproductive efficiency, including testicular development and spermatogenesis, female conception rate, and sexual differentiation, among others (Lynch and Walsh, 2007; Chang et al., 2013; Fortes et al., 2013; Lyons et al., 2014; De Camargo et al., 2015; Taylor et al., 2018; Pacheco et al., 2019). The inclusion of the effect of X chromosome markers also provided positive results by increasing the accuracy of genomic breeding value predictions (Su et al., 2014).

The evaluation and quantification of these effects is therefore extremely important to achieve expressive genetic gains and consequently efficient economic returns in production systems of Nellore cattle in Brazil, in which reproductive traits are up to 13 times more important economically than other traits (Brumatti et al., 2011). Therefore, the aim of this study was to evaluate the efficiency of inclusion and the prediction ability of the X chromosome for reproductive (occurrence of early pregnancy and age at first calving) and andrological traits (scrotal circumference) in the Nellore beef cattle herd in order to generate knowledge and applications for non-autosomal inheritance mechanisms.

## 2. Material and Methods

### 2.1 Phenotypic data

Reproductive efficiency data from male and female Nelore animals born between 1996 and 2012 were used. The animals belonged to herds in the southeastern and northeastern regions of Brazil that participate in the DeltaGen® breeding program.

The reproductive management adopted by the program since 1996 consists of two breeding seasons with a duration of approximately 70 days. An anticipated breeding season lasting approximately 60 days usually occurs between February and April, when heifers are exposed early at 14-16 months of age. All heifers are exposed to bulls regardless of their weight and body condition. Artificial insemination, controlled mating or multiple-sire mating (sire:cow ratio of 1:50) is used for this purpose. Pregnancy is confirmed approximately 60 days after the end of the anticipated breeding season. Heifers that do not conceive during this season are again exposed at 2 years of age. The criteria for culling females are reproductive failure up to 2 years of age, cow failure in conceive, and poor performance of the offspring. A small percentage is culled due to health reasons.

The occurrence of early pregnancy (P16) was defined based on the conception and calving of the heifer, given that the animal entered the breeding season at about 16 months of age. Value 2 (success) was assigned to heifers that calved at less than 31 months and value 1 (failure) to those that did not. Age at first calving (AFC), measured in days, was obtained as the difference between the date of first calving and the date of birth of the female. Scrotal circumference (SC) was measured in males at yearling.

Four seasons of birth were considered: 1 (January to March); 2 (April to June); 3 (July to September); 4 (October to December). The contemporary groups were formed using the GLM procedure of the SAS 9.2 software (SAS Institute, Inc., Cary, NC). The contemporary group for which the model showed the highest coefficient of determination for AFC and P16 was formed by the concatenation of the following information: management group at birth and yearling, season, year and farm of birth. For SC, the contemporary group comprised the management group at birth and yearling, season and farm of birth. Contemporary groups with fewer than five observations were removed from the analysis, as were records of AFC and SC outside the interval given by the mean of the contemporary group ± three standard deviations. For the analyses, a pedigree file containing the identification of the animal, sire and dam was used, totaling 99,002 animals in the relationship matrix. All available relationships (up to 11 generations) were considered. Table 1 shows the descriptive statistics of the data after the verification of consistency.

**Table 1.** Number of observations (N), mean (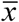), standard deviation (SD), coefficient of variation (CV), minimum, and maximum of the traits studied.

| Trait | N | $\bar{x}$ | SD | CV | Minimum | Maximum |
| --- | --- | --- | --- | --- | --- | --- |
| P16 (%) | 74,081 | 32.72* | - | - | - | - |
| AFC (days) | 75,163 | 1,027.48 | 113.97 | 10.88 | 648 | 1,200 |
| SC (cm) | 79,300 | 26.73 | 3.50 | 13.11 | 13 | 42 |

SC (scrotal circumference); AFC (age at first calving); P16 (occurrence of early pregnancy). \*Percentage of pregnant females at 16 months of age.

### 2.2 Genotypic data

For this study, 3,263 genotypes of females and males born between 1993 and 2012 were used. Performance and genealogical records are available for all genotyped animals, except for part of the sires. The animals were genotyped with the high-density Illumina BovineHD BeadChip array (Illumina Inc., San Diego, CA, USA) that contains 777,962 SNPs. The following criteria were used for quality control of the SNPs: a minor allele frequency < 0.05 and a call rate < 0.90. For annotation of X chromosome genotypes in females, value 0 was assigned to animals homozygous for the first allele (A), value 1 to heterozygous animals, and value 2 to animals homozygous for the second allele (B). For males, hemizygotes for allele A received value 0 and hemizygotes for allele B received value 2. The X chromosome containing 39,367 SNPs and autosomes were included in the analyses of the three traits, resulting in 487,490 SNPs and 3,200 animals after final cleaning data. A total of 463,753 SNPs remained for the analyses are from autosomes. Chromosomes 1 and 2, which are the two chromosomes with the largest number of SNPs in cattle and they contributed with 46,495 and 40,056 SNPs, respectively.

### 2.3 Genomic Prediction

Genomic prediction for SC, AFC and P16 was carried out considering two scenarios: 1) only autosomal markers or 2) autosomal + X chromosome markers. Analysis was performed by Bayesian inference using the GIBBS2F90 software for SC and AFC and THRGBBIS1F90 for P16 (Misztal, 2020). The phenotypes of P16, AFC and SC were used as dependent variables.

Gibbs chains of 1,000,000 cycles were generated, with a burn-in period of 50,000 cycles and a thinning interval of 50 cycles. The generated chains were analyzed with the POSTGIBBSF90 program (Misztal, 2020) to verify convergence and dependence between samples. All evaluations were performed using single-trait analysis. The model included the effects of contemporary group, genomic breeding values of the animals, and residual effect. The full model can be written in matrix notation as:

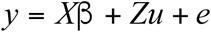

where *y* is the vector of observations for animals with or without genotype; *X* and *Z* are incidence matrices that associate β and *u* with the observations, respectively; β is the vector of contemporary group effects; *u* is the vector of direct additive genetic effects, and *e* is the vector of residual effects. A threshold model was adopted for P16, which relates the observations on a categorical scale to a normal continuous scale.

The following assumptions were established for the model: a normal prior distribution with zero mean and σ^*2*^_*a*_*H* variance was assumed for *u*, where *H* is the genomic and pedigree-based relationship matrix and σ^*2*^_*a*_ is the additive genetic variance. For the residual effects, a normal prior distribution with zero mean and *I*σ^*2*^_*e*_ variance was assumed, where *I* is the identity matrix and σ^*2*^_*e*_ is the residual variance. Since σ^*2*^_*e*_ is not estimated for the underlying scale, value 1 was attributed to σ^*2*^_*e*_ for P16. The following distribution was assumed for matrix *H* (Aguilar et al., 2010):

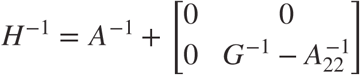

where *A* is the pedigree-based relationship matrix; *A*_*22*_ is the matrix of genotyped animals, and *G* is the genomic relationship matrix for genotyped animals estimated according to VanRaden (2008):

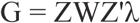

where *Z* is the matrix of coefficients adjusted for allele frequency; *D* is the matrix of weighting factors for SNPs (initially W = I), and *λ* is the weighting factor based on SNP frequencies.

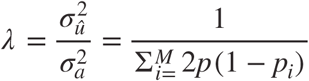

where *M* is the number of SNPs, and *p*_*i*_ is the frequency of the second allele of marker *i*.

Based on the mean estimates of the (co)variance components obtained with the model that included the X chromosome + autosomes and the model that only included autosomes, the animals were classified according to the breeding values calculated by the single-step method proposed by Aguilar et al. (2009). Unlike the traditional method that tests one marker at a time and only uses genotyped animals with a known phenotype, the single-step method combines pedigree, phenotype and genotype information of animals for which these data are available with animals that were not genotyped. Thus, two genomic breeding values were obtained for each animal and each trait, one for the model including the X chromosome (GBV_X) and one for the model including only autosomes (GBV).

To evaluate the effect of inclusion of the X chromosome on selection, the responses to the selection performed were compared including or not the X chromosome in the evaluation of the traits. Two selection scenarios were simulated, with the proportion of selected animals ranging from 1 to 10%. In the first scenario, the animals were selected based on the breeding value obtained in the analysis that did not include the X chromosome (GBV). In the second scenario, the animals were selected based on the genomic breeding values considering the X chromosome (GBV_X). The mean genomic breeding values of each model were then calculated for each trait in the selected animals. Considering that in the presence of the X chromosome the second scenario would be more adequate, the mean GBV_X were compared when selection was based on GBV and GBV_X for the three traits. The prediction ability was calculated by correlating the predicted genomic breeding values obtained by the analyses with and without the X chromosome.

## 3. Results

Higher heritability estimates (Table 2) were obtained for SC (0.40 and 0.31), AFC (0.11 and 0.09) and P16 (0.43 and 0.38) for the analyses including the X chromosome compared to those without the chromosome, respectively.

**Table 2.** (Co)variance components 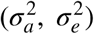 and heritabilities (*h*^2^) for scrotal circumference, age at first calving and early pregnancy of males and females in analyses including or not the X chromosome.

| Parameter | SC | SC_X | AFC | AFC_X | P16 | P16_X |
| --- | --- | --- | --- | --- | --- | --- |
| $\sigma_a^2$ | 2.68 | 3.22 | 569.67 | 668.70 | 0.07 | 0.10 |
| $\sigma_e^2$ | 5.96 | 4.45 | 5,184.45 | 5,189.34 | 0.11 | 0.13 |
| $h^2$ | 0.31 | 0.40 | 0.09 | 0.11 | 0.38 | 0.43 |

SC, SC\_X; AFC, AFC\_X; P16, P16\_X: analysis of scrotal circumference, age at first calving and early pregnancy considering only autosomes and autosomes + X chromosome, respectively.

For a better illustration, Figure 1 shows the posterior distributions of heritability for all traits considering or not the X chromosome. Observation of the graph shows that, based on the intersection of densities, inclusion of the X chromosome exerted a greater effect on the difference between estimates for SC and P16 than for AFC. Despite the smaller difference in AFC, in terms of reproductive trait, this difference is still considerable.

**Figure 1.**
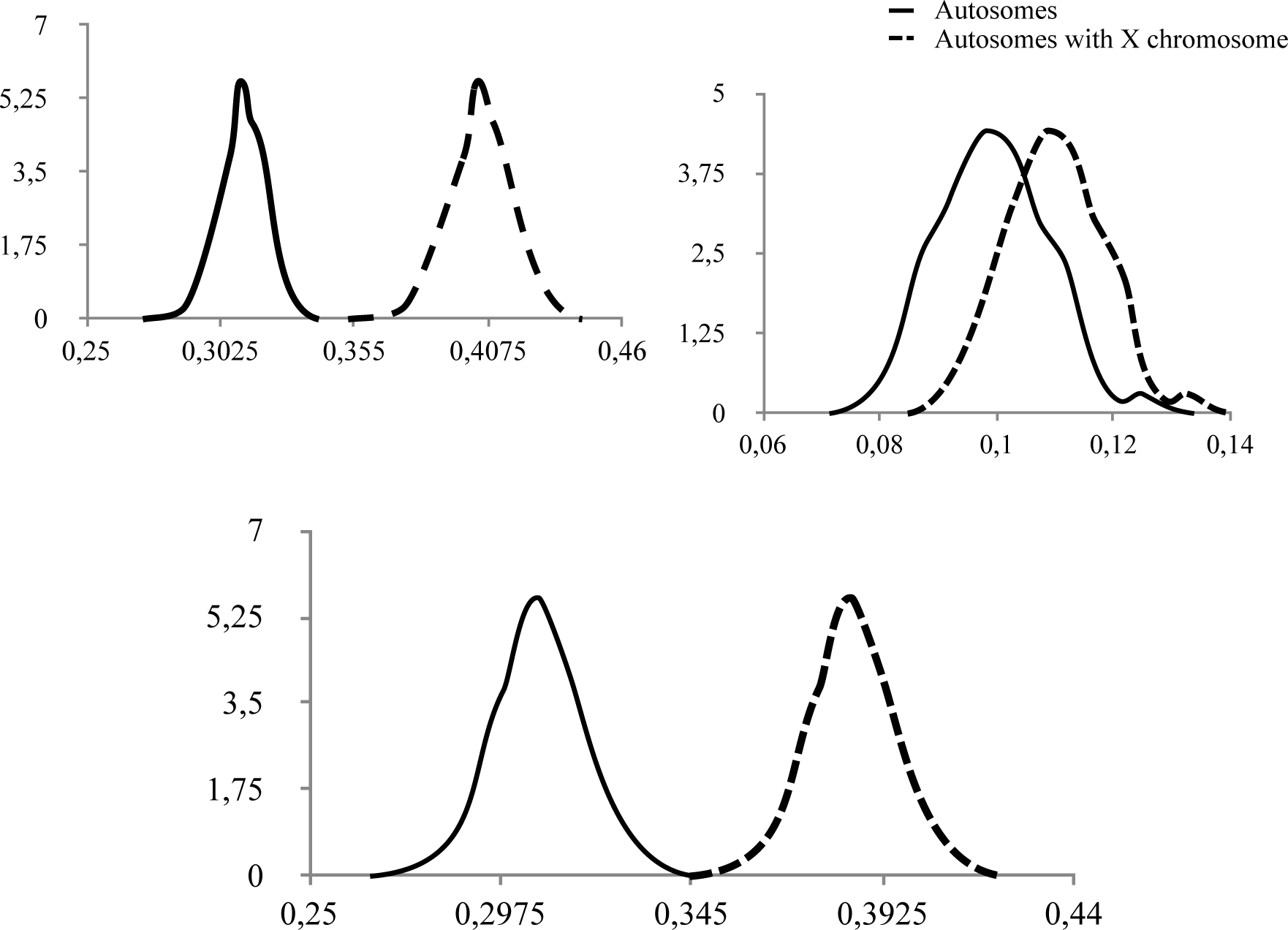
Posterior densities of the heritability (h^2^) for scrotal circumference (SC), age at first calving (AFC), and early pregnancy (P16) considering or not the X chromosome.

Table 3 shows the mean genomic breeding values of the selected animals obtained in analyses that included the X chromosome and the percent reduction in these mean values when selection is based on the results of analysis that did not include the X chromosome according to the selected fraction (GBV_X_SC - 16.51; 14.233; 13.45; GBV_X_AFC – 65.01; 32.73; 20.32 and GBV_X_P16 – 0.56; 0.33; 0.19, for 1%, 5% and 10% of selected fraction). For SC, when the effects of the X chromosome are ignored, the loss can reach slightly more than 7% of the mean genomic breeding value of the selected animals. For P16, the loss can reach more than 4%, while this loss does not seem to be as important for AFC.

**Table 3.** Mean genomic breeding values for analyses including the sex chromosomes when 1, 5 and 10% of the top males are selected and percent reduction (in parentheses) when selection is based on the analysis that considers only autosomes.

| P(%) | GBV_X_SC | GBV_X_AFC | GBV_X_P16 |
| --- | --- | --- | --- |
| 1 | 16.51 | 65.01 | 0.56 |
|  | (-7.91%) | (-1.91%) | (-4.11%) |
| 5 | 14.23 | 32.73 | 0.33 |
|  | (-6.61%) | (-2.87%) | (-3.43%) |
| 10 | 13.45 | 20.32 | 0.19 |
|  | (-5.12%) | (-2.48%) | (-3.06%) |

P(%): percentage of selected animals; GBV\_X\_SC, GBV\_X\_AFC and GBV\_X\_P16: mean genomic breeding values of animals for scrotal circumference, age at first calving and early pregnancy, respectively.

Greater differences between the predicted genomic breeding values were observed for P16 and SC in situations when the selection intensity was higher (1 and 5%). When comparing models that fit SNPs using autosomes versus autosomes + X chromosome, a question that may arise is whether the predictive performance is improved due to the biological role of the SNP on the X chromosome or simply because the model is fitting a larger set of SNPs. To answer this question, 50 different and random sets of 39k SNPs of the autosomes (SNP density of the X chromosome in the study) were removed and the prediction ability of these sets was evaluated (Table 4). The mean estimates obtained did not change for the 50 replicates, producing an average predictive correlation of 0.79, 0.98 and 0.84 for SC, AFC and P16, respectively.

**Table 4.** Pearson’s correlation coefficients between predicted genomic breeding values of animals for scrotal circumference (SC), age at first calving (AFC), and early pregnancy (P16) obtained in analyses that included or not the X chromosome.

|  | GBV_SC | GBV_AFC | GBV_P16 |
| --- | --- | --- | --- |
| GBV_XSC | 0.79 | -- | -- |
| GBV_XAFC | -- | 0.98 | -- |
| GBV_XP16 | -- | -- | 0.84 |

GBV\_\*: genomic breeding value predicted in analyses including only autosomes; GBV\_X\*: genomic breeding values predicted in analyses including the X chromosome.

For SC and P16, the prediction ability of the breeding values given by Pearson’s correlation (Table 4) shows a greater difference when compared to AFC; hence, greater care in the ranking of animals is necessary for these two traits when the X chromosome and males are not included in the analyses.

## 4. Discussion

The estimates of heritabilities (Table 2) agree with those reported in the literature for the same traits (Lemos et al., 2015; Sousa et al., 2015; Ambrosini et al., 2016; Buzanskas et al., 2017; Carvalho et al., 2019). Reproductive efficiency traits are largely affected by the environment and by the non-additive genetic fraction. The increase in the heritability estimates can be explained by the fact that the inclusion of the X chromosome reduced the noise caused by the non-additive fraction, with a consequently better fit of the effects. The larger number of genotypes considered in the analysis permits to improve and to separate additive and dominance effects that influence the phenotypes, thus improving estimations.

Another explanation is that inclusion of the X chromosome accounts for additive effects and the interaction of its genes. The X chromosome harbors three times more genes that are expressed in the female reproductive system and 15 times more genes related to male germ cells than autosomes (Liu, 2019). This fact highlights the importance of considering the additivity of X chromosome genes. By not including X chromosome markers, the effect of autosomal markers is inflated (data not shown). Using the WssGBLUP method, Irano et al. (2016) emphasized that, when an iterative procedure for updating SNP effects is used, adjustment of the effects tends to be more accurate at each iteration since the combination of weights minimizes prediction errors and reflects the reality of adjacent SNPs that contribute to the expression of the trait. However, if SNPs present on the X chromosome are not accounted for and these markers have important pleiotropic effects, false-positives may be accentuated. Furthermore, X chromosome genes can epistatically interact with one another and with autosomal genes. Epistasis is partially heritable, a fact that helps explain the increase in additive genetic variance. Accordingly, Su et al. (2014), who evaluated genomic predictions in analyses that included or not the X chromosome, concluded that the model considering the polygenic effect presented a loss of accuracy in the estimates by excluding X chromosome markers. Pacheto et al. (2019) also obtained better genomic predictions for bull fertility when markers in regions of the X chromosome were considered.

The observation of different heritabilities for the same trait depending on the model used suggests that the genotypes react differently to inclusion of the X chromosome, reinforcing the existence of gene interactions. Fortes et al. (2012), analyzing transcription networks, demonstrated important effects of gene interactions between the X chromosome and autosomes on reproductive traits in males and females, in agreement with the results obtained in this study.

Although the different heritability estimates indicate divergence in the genotype performance of the animals in the different analyses of the three traits, this difference only has consequences from the practical point of view of selection when it causes changes in the relative merit of the animals. Once this difference in the estimates became evident, it was important to quantify the effect of this relationship on the selection of animals by comparing the analysis including only autosomal genes with that considering the presence of the X chromosome (Tables 3 and 4), since most of the currently performed evaluations do not include this chromosome. For AFC the loss of percent reduction from mean genomic breeding values does not seem to be as important as for SC and P16 traits. Within this context, Carvalho et al. (2019), who worked with part of the dataset used here, showed that the X chromosome has little influence on AFC in Nelore cows. This might be a particularity of the trait and/or population, or which may also be related to the presence of variance heterogeneity for the two traits (SC and P16). Pacheco et al (2019), evaluating the effect of sex chromosomes on reproductive traits in males and females, found an increase of 7.5% in the prediction ability of a model that considered the X chromosome together with the autosomes.

The prediction ability of the 50 different and random sets of 39k SNPs (Table 4), notably, shows that the differences in predictions including X chromosome markers are due to their biological role and not to the increased number of markers. These estimates demonstrate that inclusion of the X chromosome in the analysis can improve the prediction of genomic breeding values, especially for SC. Consequently, better genetic gain can be obtained for selection based on these evaluations. These results suggest that the X chromosome must not be overlooked when the aim is to fully understand the genetic basis of reproductive efficiency traits.

Analyses that include the X chromosome and all animals (males and females) provide more reliable genomic breeding value estimates by considering both genetic and environmental differences for the three traits studied, thus reducing estimation bias. However, it is important to note that reproductive efficiency traits have a highly complex genetic architecture and further studies involving different populations are therefore necessary. There is still a need to improve the genomic selection results for reproductive traits in order to increase the understanding of specific genes and of polymorphisms underlying genetic variations at the population level. This goal has been achieved in relatively few cases so far; however, there are still opportunities for improvements in this area.

## 5. Conclusion

Considering the present results and the fact that the use of a model that does not include the effect of X chromosome markers led to considerable losses, the evaluation of reproductive traits should combine X chromosome and autosomal markers.

## Acknowledgements

Funding: This work was supported by the National Council for Scientific and Technological [CNPq grant numbers 422799/2016-5]; São Paulo Research Fundation [FAPESP grant numbers 2009/16118-5].

## Author Contributions Statement

In this study, the co-authors: Iara Del Pilar Solar Diaz, Gregório Miguel Ferreira de Camargo and Valdecy Aparecida Rocha da Cruz performed the genetics analysis as the causal inference of the full manuscript. Isis da Costa Hermisdorff, Caio Victor Damasceno Carvalho, Lucia Galvão de Albuquerque, Raphael Bermal Costa participated in substantial contribution to conception and revising it critically for important intellectual content. All the authors in this manuscript have read and approved the final version.

